# Paecilomycone inhibits quorum sensing in Gram-negative bacteria

**DOI:** 10.1101/2022.09.12.507720

**Authors:** Wouter A. G. Beenker, Jelmer Hoeksma, Marie Bannier-Hélaouët, Hans Clevers, Jeroen den Hertog

## Abstract

*Pseudomonas aeruginosa* is an opportunistic pathogen that causes major healthcare concerns due to its virulence and high intrinsic resistance to antimicrobial agents. Therefore, new treatments are highly needed. An interesting approach is to target quorum sensing (QS). QS regulates the production of a wide variety of virulence factors and biofilm formation in *P. aeruginosa*. This study describes the identification of paecilomycone as inhibitor of QS in both *C. violaceum* and *P. aeruginosa*. Paecilomycone strongly inhibited the production of virulence factors, including various phenazines, and biofilm formation. In search of the working mechanism, we found that paecilomycone inhibited the production of 4-hydroxy-2-heptylquinoline (HHQ) and 3,4- dihydroxy-2-heptylquinoline (PQS), but not 2’-aminoacetophenone (2-AA). We suggest that paecilomycone affects QS in *P. aeruginosa* by targeting the PqsBC complex and alternative targets, or alters processes that influence the enzymatic activity of the PqsBC complex. The toxicity of paecilomycone towards eukaryotic cells and organisms was low, making it an interesting lead for further clinical research.

**Importance:** Antibiotics are becoming less effective against bacterial infections due to the evolution of resistance among bacteria. *Pseudomonas aeruginosa* is a Gram-negative pathogen that causes major healthcare concerns and is difficult to treat due to its high intrinsic resistance to antimicrobial agents. Therefore, new targets are needed and an interesting approach is to target quorum sensing (QS). QS is the communication system in bacteria that regulates multiple pathways including the production of virulence factors and biofilm formation, which leads to high toxicity in the host and low sensitivity to antibiotics, respectively. We found a compound, named paecilomycone, which inhibited biofilm formation and the production of various virulence factors in *P. aeruginosa.* The toxicity of paecilomycone towards eukaryotic cells and organisms was low, making it an interesting lead for further clinical research.

## Introduction

*Pseudomonas aeruginosa* is a Gram-negative pathogen that causes nosocomial infections in immunocompromised patients. It is involved in a variety of acute and chronic infections including urinary tract infections, burns or wound infections, and in respiratory diseases like cystic fibrosis (CF)(1, 2). *P. aeruginosa* has a high intrinsic resistance due to the low permeability of its outer membrane, the high amount of efflux pumps and the capability to form biofilms. Therefore, *P. aeruginosa* infections are difficult to treat(1, 3). Once established in the lung, *P. aeruginosa* infections often become chronic and 60-80% of adult CF patients harbor chronic *P. aeruginosa* infections, contributing to morbidity and mortality of the patients(4, 5). Therefore, new treatments against *P. aeruginosa* infections are highly needed, and quorum sensing (QS) as target might be an interesting approach.

QS in bacteria is a communication system that involves changes in gene expression in response to cell density. In Gram-negative bacteria, the QS systems are very similar with small modifications. These systems contain homologues of a LuxI-type synthase that produces acylated homoserine lactones (AHL) that are specific to that bacterial strain. When the number of bacteria increases, the concentration of AHL increases. The AHL will bind to its cognate LuxR-type receptor, which subsequently binds to the promoter of QS target genes and alters gene expression(6–8).

In *P. aeruginosa* there are three different QS systems. Two of these systems are typical Gram-negative bacterial LuxI/R-type QS systems, the *las*-encoded system and the *rhl*-encoded system. The third system in *P. aeruginosa* is a unique system based on 4-hydroxy-2-alkylquinolines (HAQs) that are synthesized by the enzymes of the *pqsABCDE* operon and PqsH. Among these HAQs are 3,4-dihydroxy-2-heptylquinoline (PQS) and its precursor 4-hydroxy-2-heptylquinoline (HHQ) that can bind to PqsR (also called MvfR)(9–11). These three QS systems in *P. aeruginosa* play a major role in the production of virulence factors and biofilm formation. Therefore, inhibition of QS leads to decreased production of virulence factors, including pyocyanin, rhamnolipids, and elastase and a concomitant decrease in biofilm formation(12, 13). This way, QS inhibitors might lead to reduced toxicity of the bacteria towards the host and at the same time higher susceptibility to antibiotics(14, 15).

In a previous study from our lab, we reported the search for novel QS inhibitors among secondary metabolites of 10,207 strains of fungi(16). Here, we report identification of paecilomycone from *Aspergillus allahabadii* as QS inhibitor in *C. violaceum* and *P. aeruginosa*. Paecilomycone treatment showed a great reduction in biofilm formation, phenazine production and HAQ synthesis. In addition, toxic effects appeared to be low, especially in complex systems, making paecilomycone an interesting molecule for further development for clinical use.

## Materials and methods

### Bacterial strains and growth conditions

Bacterial strains used in this study (Supplementary Table 1) were stored at -80 °C in 20 % glycerol stock solutions. *C. violaceum* was plated on tryptic soy agar (TSA) and grown in tryptic soy broth (TSB) at 27 °C. PAO1 strains were grown on Luria agar (LA) plates at 37 °C and grown in medium specific for the assay. *E. coli* RHO3 strains were used for conjugation and medium was supplemented with 400 µg/mL 2,6- Diaminopimelic acid (Sigma-Aldrich, Merck Life Science, Amsterdam, the Netherlands) to support growth.

Mutants were generated using allelic exchange following the method described before(17). For the generation of mutants we inserted upstream (UP.Fw and UP.Rv primers used) and downstream (DN.Fw and DN.Rv primers used) regions of the gene of interest in pEX18Gm vector (gift from Joe Harrison, University of Calgary) using Gibson assembly restriction cloning, using the restriction enzymes SacI and SphI (Supplementary Table 2 and 3). After Gibson assembly, the vector containing the regions flanking the gene of interest-was transformed into RHO3 *E. coli* donor strains before conjugation with WT PAO1 cells. Mutant cells were identified with colony PCR (seq.Fw and seq.Rv primers used) and confirmed by sequencing (performed by Macrogen Europe BV).

### Chemical analysis

#### Purification of paecilomycone

Paecilomycone was purified using the same method as described before with minor changes(16). In brief, the fungal strain *Aspergillus allahabadii* was grown on a malt extract agar (MEA) plate at 25 °C. After 7 d, cubes of 5x5 mm were cut out and 2 cubes were inoculated per 100 mL bottle containing 50 mL of potato dextrose broth (PDB). The liquid culture was inoculated at 25 °C with 100 rpm orbital shaking. After 7 d, the liquid medium was filter sterilized using a 0.22 µm Millipore filter (Merck, Amsterdam, the Netherlands). The sterile supernatant was extracted using 3x 1/3 volume of ethyl acetate and evaporated to dryness using a rotary evaporator with a water bath at 40 °C. The dried pellet was dissolved in DMSO.

The extract was fractionated using a preparative high-performance liquid chromatography (HPLC) system consisting of a Shimadzu CBM-20A controller, a Shimadzu LC-20AP pump and a Shimadzu FRC-10A fraction collector using a C18 reversed-phase Reprosil column (10 µm, 120 Å, 250 x 22 mm) and a Shimadzu SPD- 20A UV-detector set at 214 nm and 254 nm. The mobile phase consisted of 100 % MQ with 0.1 % trifluoroacetic acid (Buffer A) and 100 % acetonitrile with 0.1 % trifluoroacetic acid (Buffer B). Protocol consisted of 5 % Buffer B for 5 min, followed by a linear gradient to 95 % buffer B for 40 min, 5 min of 95 % buffer B before returning to 5 % buffer B for another 5 min with a constant flow rate of 12.5 mL/min. The active fraction with a retention time of 31 min was collected and dried overnight using a speedvac.

Next day, the dried pellet was dissolved in DMSO and fractionated again using the same preparative HPLC system but with different buffers. Buffer A consisted of 95:5 MQ:acetonitrile + 10 mM NH_4_OAc. Buffer B consisted of 80:20 acetonitrile:MQ +10 mM NH_4_OAc. Protocol consisted of 0 % Buffer B for 5 min, followed by a linear gradient to 100 % buffer B for 40 min, 5 min of 100 % buffer B before returning to 0 % buffer B for another 5 min with a constant flow rate of 12.5 mL/min. Paecilomycone had a retention time of 23 min and was collected and dried using a speedvac.

Paecilomycone was dissolved in a stock concentration of 40 mM in DMSO and stored at -20°C. Purity was checked using an analytical HPLC Shimadzu LC-2030 system with PDA detection (190-800 nm) with a Shimadzu Shim-pack GIST C18-HP reversed-phase column (3 µm, 4.6x100 mm). The mobile phase consisted of 100 % MQ with 0.1 % trifluoroacetic acid (Buffer A) and 100 % acetonitrile with 0.1 % trifluoroacetic acid (Buffer B). Protocol consisted of a linear gradient from 5 % to 95 % buffer B for 10 min, followed by 2.5 min of 95 % buffer B before returning to 5% buffer B with a constant flow rate of 1 mL/min.

#### Identification of paecilomycone

The UV-VIS spectrum of paecilomycone was obtained using a Shimadzu LC-2030C 3D Plus analytical HPLC system with PDA detection (190-800 nm) with a Reprosil-PUR 120 C18 AQ column (3 µm, 120 Å, 4.6x100 mm). The compound was further analyzed by measuring the mass using a Shimadzu LCMS-2020 system. For LC-MS analysis, the mobile phase consisted of 100 % MQ with 0.05 % formic acid (Buffer A) and 100 % acetonitrile with 0.05 % formic acid (Buffer B). Protocol consisted of a linear gradient from 5 % to 95 % buffer B for 10 min, followed by 2.5 min of 95 % buffer B before returning to 5 % buffer B with a constant flow rate of 0.5 mL/min. Paecilomycone A had a retention time of 12.7 min, paecilomycone B had a retention time of 9.8 min, and paecilomycone C had a retention time of 8.9 min.

This was followed by a more accurate high resolution mass spectrometry, for which the sample was injected followed by an injection of sodium formate for detection of sodium adduct ions. This procedure gave an internal calibrant to facilitate a more accurate measurement of the mass of the sample.

For Nuclear magnetic resonance (NMR) analysis, paecilomycone was dissolved in DMSO-d6. The NMR measurements (^1^H, ^13^C, Heteronuclear Single-Quantum Correlation (HSQC) and Heteronuclear Multiple-Bond Correlation spectroscopy (HMBC)) were performed on a Bruker 600 MHz.

### Quorum sensing inhibition in *C. violaceum*

To measure the effect on QS inhibition in *C. violaceum,* we used a protocol based on previous work with minor changes(18). In brief, *C. violaceum* was plated on tryptic soy agar (TSA) and grown in TSB overnight at 27 °C. Next morning, bacteria were diluted and grown until OD_600_ = 0.5-0.7. Then, bacteria were diluted 1000 x and added to a 96-well plate containing paecilomycone in serial dilutions up to a volume of 100 µL (range 31 nM – 1 mM). High concentrations of DMSO are toxic to the bacteria, and therefore the maximum concentration of DMSO was limited to 2.5 %. Plates were incubated for 20 h at 27 °C while shaking at 180 rpm.

To measure violacein production, the plate was centrifuged for 10 min at 3000 rpm to pellet the violacein. Subsequently, the supernatant was discarded and the violacein pellet was dissolved in 200 µL of 96 % ethanol. The plate was centrifuged for 10 min at 3000 rpm to avoid interference of cell turbidity in absorbance measurements. Half of the supernatant was then transferred to a new 96-well plate and violacein was quantified by measured the OD at 562 nm on the ASYS expert plus microplate reader (Biochrom Ltd, Cambridge, UK).

To measure the effect of paecilomycone on viability of the bacteria, resazurin staining was used. In parallel with the measurements of violacein production, another plate with *C. violaceum* and paecilomycone dilutions was prepared and incubated overnight. Next morning, the plate was centrifuged for 10 min at 3000 rpm to pellet the bacterial cells. The supernatant was aspirated and 0.1 mM resazurin (in PBS) solution was added. Plates were incubated for another 45 min at 27 °C before fluorescence was measured on a PHERAstar microplate reader (BMG Labtech) using 540 nm exication and 590 nm emission wavelength.

### Quorum sensing inhibition in P. aeruginosa

The experiments were performed as previously described(19). In brief, *P. aeruginosa* PAO1 reporter lines were grown overnight in AB minimal medium supplemented with 0.5 % glucose and 0.5 % casamino acids. Cultures were diluted until OD_450_ = 0.1-0.2 before adding to a 96-well plate containing serial dilutions of paecilomycone up to a volume of 200 µL (range 4 nM – 500 µM). The GFP fluorescence (excitation 485 nm, emission 535 nm) and absorbance (600 nm) were measured every 15 min for 15 h at 34 °C on a CLARIOstar microplate reader (BMG Labtech). IC_50_ values were calculated using PRISM software, plotting the maximum slope of GFP/OD_600_.

### Biofilm assay

Bacteria were grown overnight in AB minimal medium supplemented with 0.5 % glucose and 0.5 % casamino acids before diluting 1000 x. Diluted bacterial cells were added to a 96-well plate containing paecilomycone in serial dilutions (range 3.9 µM – 500 µM) in triplicates to a final volume of 200 µL. Plates were sealed with BreatheEasy seal (Sigma-Aldrich, Merck Life Science, Amsterdam, the Netherlands) to prevent evaporation and incubated at 37 °C under static conditions for 24 h. Next day, the medium was removed and the wells were rinsed with PBS. Biomass was stained with 0.1 % (w:v) crystal violet solution for 5 min. Crystal violet was discarded and excess crystal violet was removed by rinsing with water. Plates were dried and bound crystal violet was dissolved in 33 % (v:v) acetic acid and quantified at 562 nm using an ASYS expert plus microplate reader (Biochrom Ltd, Cambridge, UK).

### Virulence factor assays

#### Pyocyanin assay

The pyocyanin assay was performed as previously described with minor modifications(20). Bacteria were grown overnight in King’s A medium (2 % (w:v) protease peptone, 1 % (w:v) potassium sulfate, 0,164 % (w:v) magnesium chloride, 1 % (v:v) glycerol in MQ), before diluting them 100 x. The assay was performed in bacterial tubes containing 1 mL of diluted bacterial culture in combination with paecilomycone at desired concentrations (range 2 µM – 125 µM). A Δ*lasI*/Δ*rhlI* QS mutant strain was included as negative control. The tubes were incubated at 37 °C on an orbital shaker set at 180 rpm. After 24 h, bacterial cells were pelleted by centrifugation at 4000 rpm for 10 min and 900 µL supernatant was extracted with a similar volume of chloroform. 800 µL of chloroform was then added to 700 µL of 0.2 M HCl and samples were mixed well. The HCl phase was then measured at 520 nm to measure relative pyocyanin concentrations.

#### Rhamnolipid assay

The rhamnolipid assay was performed as previously described with minor modifications(21). Bacteria were grown overnight in AB minimal medium supplemented with 0.5 % glucose and 0.5 % casamino acids, before diluting 100 x. The assay was performed in bacterial tubes containing 1 mL of diluted bacterial culture in combination with paecilomycone at desired concentration (range 2 µM – 125 µM). A Δ*lasI*/Δ*rhlI* QS mutant strain was included as negative control. The tubes were incubated at 37 °C on an orbital shaker set at 180 rpm. After 24 h, bacterial cells were pelleted and 900 µL of supernatant was added to diethyl ether (1:1, v:v) and tubes were shaken vigorously. The diethyl ether layer was then transferred to a fresh tube and dried at RT. 100 µL was added to dissolve the dried pellet before addition of 800 µL of 12.9 mM orcinol (Sigma Aldrich, Merck Life Science, Amsterdam, the Netherlands) in 70 % (v:v) H_2_SO_4_. The reaction was maintained at 80 °C for 30 min before measuring the absorbance at 495 nm.

#### Phenazine and HAQ assay

The phenazines and HAQ assay was based on the pyocyanin assay with minor modifications. Bacteria were grown in King’s A medium before 100 x dilution. The assay was performed in bacterial tubes containing 3 mL of diluted bacterial culture in combination with paecilomycone at desired concentrations (range 1 µM – 125 µM). A Δ*lasI*/Δ*rhlI* QS mutant strain was included as negative control. The tubes were incubated at 37 °C with 180 rpm orbital shaking. After 24 h, the bacterial cells were pelleted and the supernatant was extracted with 2 x 1 mL chloroform. Chloroform was dried overnight and the pellet was dissolved in DMSO. Extracts were run on previously described analytical HPLC and compounds were quantified by using calibration curves made by serial dilutions of standard commercial phenazines and HAQs. If needed, extra verification of the compounds was performed using LC-MS.

### Toxicity assays

#### HepG2 toxicity assay

HepG2 cells were seeded in 96-well plates and grown in DMEM low glucose medium (ThermoFisher Scientific, 10567014) supplemented with 10 % FBS. Cells were grown until a confluence of approximately 70-80 % before addition of paecilomycone (range 244 nM – 500 µM) in triplicates, with a final DMSO concentration of 1 % DMSO. Treated cells were incubated at 37°C with 5 % CO_2_ for 24 h. To measure the viability of the cells, 0.1 mM resazurin (Sigma-Aldrich) solution was added and cells were incubated for another 3 h. Fluorescence intensity was measured on a PHERAstar microplate reader (BMG Labtech), using an excitation wavelength of 540 nm and emission wavelength of 590 nm.

#### Organoid toxicity assay

Human colon tissue was obtained from the UMC Utrecht with patient informed consent. The patient was a male diagnosed with small colon adenocarcinoma. After resection, a sample from non-transformed, normal mucosa was taken for this study. This study was approved by the UMC Utrecht (Utrecht, the Netherlands) ethical committee and was in accordance with the Declaration of Helsinki and according to Dutch law. This study is compliant with all relevant ethical regulations regarding research involving human participants. All organoids experiments were performed in the lab of Hans Clevers.

Human colon organoids were maintained in human colon expansion medium as previously described(22). The toxicity assay was largely performed as described elsewhere(23). In brief, 2 d after the previous split, organoids were dissociated from the Cultrex Basement Membrane Extract (BME, R&D Biosystems, Bio-Techne, 3533- 001-02) using dispase (ThermoFisher Scientific, 17105-041) for 30 min at 37 °C. Then, organoids were washed in advanced DMEM/F12 (ThermoFisher Scientific, 12634- 010) supplemented with penicillin-streptomycin (ThermoFisher Scientific, 15140122), Hepes (ThermoFisher Scientific, 15630080) and GlutaMAX (ThermoFisher Scientific, 35050061) [hereafter called washing medium], filtered through a 70 µM cell strainer (Greiner) and pelleted at 500 g for 5 min. The pellet was resuspended in 1 mL of washing medium and organoids were counted. Organoids were resuspended at a concentration of 18,750 organoids/mL in 5 % cold BME/95 % cold human colon expansion medium. 40 µL of organoid suspension (750 organoids) was dispensed in each well of a 384 well-plate (Corning, 4588) using a multi-drop combi reagent dispenser (ThermoFisher Scientific, 5840300). Paecilomycone was added immediately after plating the organoids at concentrations ranging from 10 nM to 200 µM using the Tecan D300e Digital dispenser (Tecan). The stock concentration was 10 mM and hence, the highest concentration contained 2 % DMSO. Therefore, a 2 % DMSO only viability control was added. Organoids were incubated in a humidified incubator with 5 % CO_2_ at 37 °C for 5 d. After 5 d, the ATP levels were measured by a Cell Titer GLO 3D assay (Promega, G9681) following the manufacturer’s instructions. Luminescence was measured using a Spark multimode microplate reader (Tecan) with an integration time of 500 ms. Results were normalized to 1 % DMSO control (100 % viability) and 1 µM staurosporine (0 % viability). Concentrations were measured in triplicates.

#### Zebrafish toxicity assay

Zebrafish eggs were obtained from Tubingen long fin family crosses. The zebrafish embryos were allowed to develop normally for 48 h. At 48 hpf the embryos were divided over 24-well plates, 10 embryos per well in 1 mL of fresh E3-medium. Subsequently, paecilomycone was added in serial dilutions to the wells (range 3.9 nM – 250 µM). The effects were scored at 72 hpf.

All procedures involving experimental animals were approved by the local animal experiments committee (Koninklijke Nederlandse Akademie van Wetenschappen- Dierexperimentencommissie) and performed according to local guidelines and policies in compliance with national and European law. Adult zebrafish were maintained as previously described(24).

### Commercial compounds used

2’-aminoacetophenone (2-AA), 4-hydroxy-2-heptylquinoline (HHQ), 2-heptyl-3- hydroxy-4(1H)-quinolone (PQS) (Sigma Aldrich, Merck Life Science, Amsterdam, the Netherlands), 2-heptyl-4-quinolinol 1-oxide (HQNO), 1-phenazinecarboxylic acid (PCA), and phenazine-1-carboxamide (PCN) (Caymen Chemicals), used to make calibration curves to quantify molecules in bacterial extract. Funalenone (AdipoGen Life Sciences) to compare activity with paecilomycone.

## Results

### Identification of paecilomycone

For the purification of paecilomycone, we grew the fungus *Aspergillus allahabadii* in PDB at 25°C with 100 rpm shaking. After 7 days, the liquid culture was filter-sterilized, to separate the fungus from the secondary metabolites. Organic compounds were isolated by liquid-liquid extraction, dried using a rotary shaker and the pellet was dissolved in DMSO. The extract was then fractionated using a preparative high-performance liquid chromatography (HPLC) system, using trifluoroacetic acid (TFA) as modifier (Supplementary Figure 1A). The fractions were tested for quorum sensing (QS) inhibitor activity using *C. violaceum* as reporter bacteria and violacein production as read-out. *C. violaceum* has a single LuxI/R QS network (CviI/R) and produces a purple pigment, named violacein, upon activation of this QS network(25). This makes this bacterium an easy reporter to search for novel QS inhibitors(18). Fraction 20 showed inhibition of violacein production.

To purify the active compound, the active fraction was fractionated again on aforementioned preparative HPLC using ammonium acetate as modifier to be able to further purify the fraction (Supplementary Figure 1B). Again, the fractions were tested, and fraction 1 showed QS inhibitor activity.

When running fraction 1 on analytical HPLC, we found three peaks (Figure 1A). Peak 1 was the largest and showed a characteristic UV-VIS spectrum (**213**(**100**), 238sh, 277sh, 388(40)) and a m/z of 289.1 [M+H] and 287.0 [M-H] suggesting a nominal mass of 288 (Figure 1B, E, H). In addition, high-resolution mass spectrometry measured a mass of 289.5937, giving a calculated mono-isotopic mass of 289.0692 [M+H], with a molecular formula prediction of C_15_H_12_O_6_ (Supplementary Figure 2). This UV-VIS spectrum and mass showed similarities with paecilomycone A(26). Therefore, the NMR data were compared to the published data of paecilomycone A (Table 1, Supplementary Figure 3).

**Figure 1:**
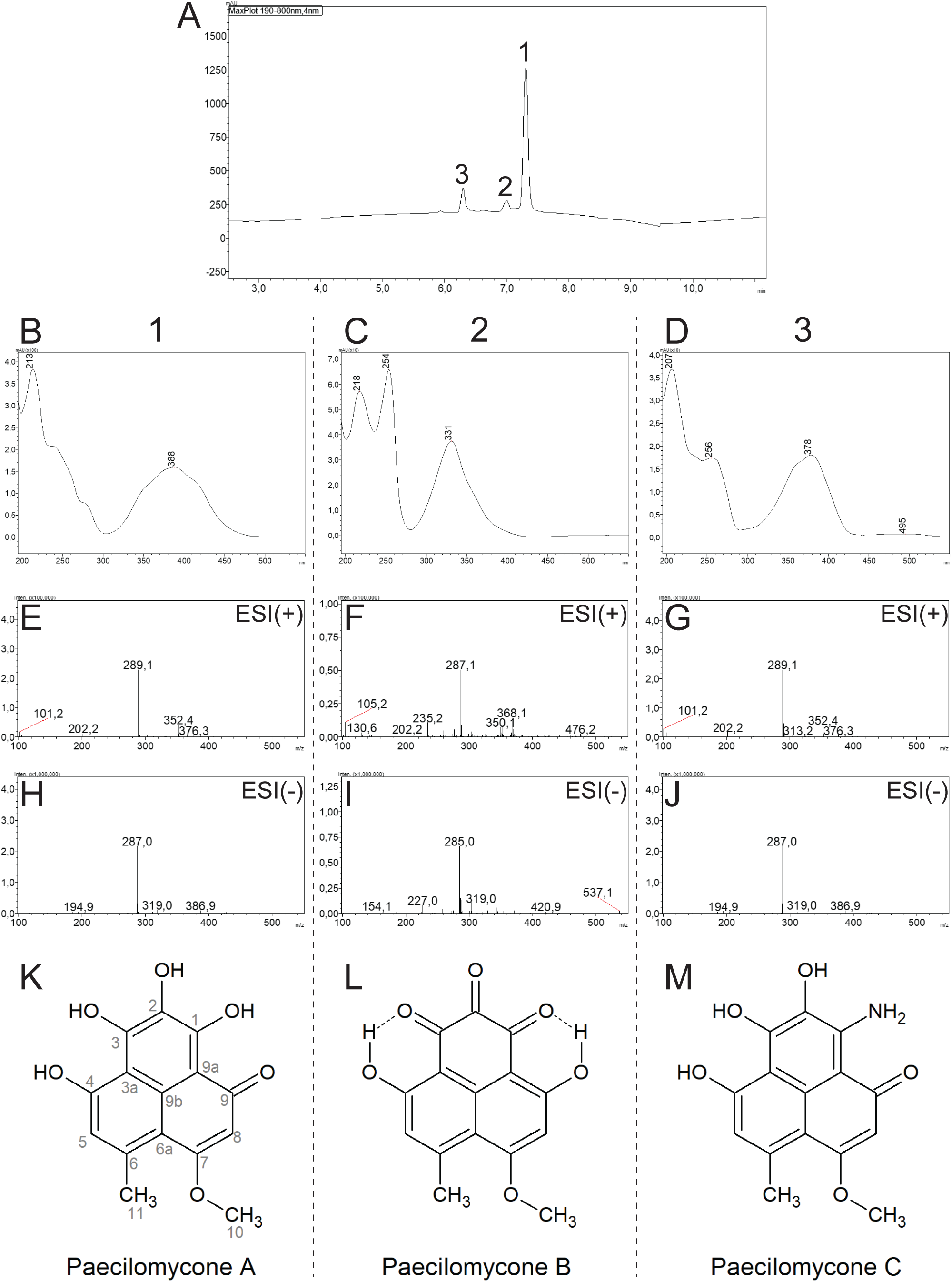
Identification of paecilomycone. A) aHPLC spectrogram of paecilomycone showing 3 peaks. UV-VIS chromatogram of B) peak 1, C) peak 2, D) peak 3. LC-MS ESI^+^ (positive mode) spectrogram of E) peak 1, F) peak 2, G) peak 3. LC-MS ESI^-^ (negative mode) spectrogram of H) peak 1, I) peak 2, J) peak 3. Chemical structures of paecilomycones belonging to K) peak 1, L) peak 2, M) peak 3.

**Table 1.**
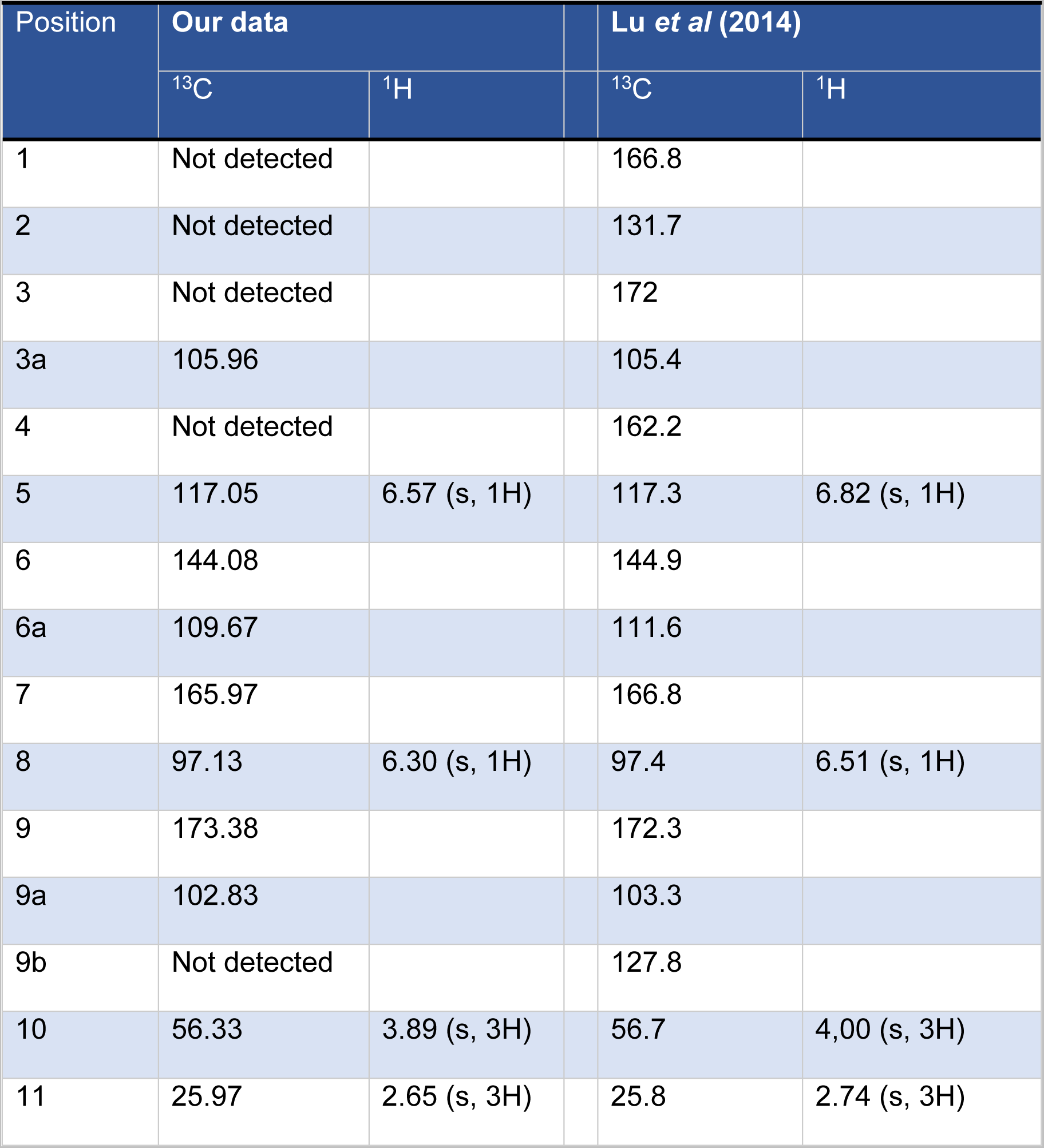
^13^C and ^1^H NMR spectral data of paecilomycone compared to Lu *et al* (2014)^31^ in DMSO-*d_6_*

Whereas not all carbons were detected, the NMR data, like the UV-VIS and LC-MS, showed high similarities with the data of Lu *et al.*(26). For further validation, the HMBC data were compared (Supplementary Figure 4). The HMBC data showed the same correlations as described before by Lu *et al.*(26), validating that this compound is indeed paecilomycone A (Figure 1K).

However, next to paecilomycone A, two other small peaks were observed in the analytical HPLC chromatogram (Figure 1). Peak 2 showed a UV-VIS spectrum (218(90), **254**(**100**), 331(50)) and a m/z of 287.1 [M+H] and 285.0 [M-H] suggesting a nominal mass of 286, corresponding to paecilomycone B (Figure 1C, F, I, L)(26). Peak 3 showed a UV-VIS (**207**(**100**), 259 (50), 378 (50)) and a m/z of 289.1 [M+H] and 287.0 [M-H] suggesting a nominal mass of 288, corresponding to paecilomycone C (Figure 1D, G, J, M)(26). The ratio of the various paecilomycones was approximately 85:5:10 (paecilomycone A: B: C, respectively), when dissolved in acetonitrile/water with 0.1% TFA. Using this set-up it was not possible to separate the various paecilomycones further. Therefore, the combination of the three paecilomycones will be referred to as paecilomycone, unless stated otherwise. In conclusion, the fraction of *Aspergillus allahabadii* with QS inhibitor activity contained paecilomycone.

### Purified paecilomycone shows QS inhibitor activity in *C. violaceum*

Purified paecilomycone strongly inhibited violacein production in *C. violaceum* with an IC_50_ of 72.5 µM (Figure 2A). Viability was measured in parallel to distinguish between effects on QS and effects on bacterial growth that could affect violacein production. No toxic effects of paecilomycone were detected on bacteria at these concentrations. Therefore, paecilomycone was a potent QS inhibitor at concentrations that do not affect cell viability.

**Figure 2:**
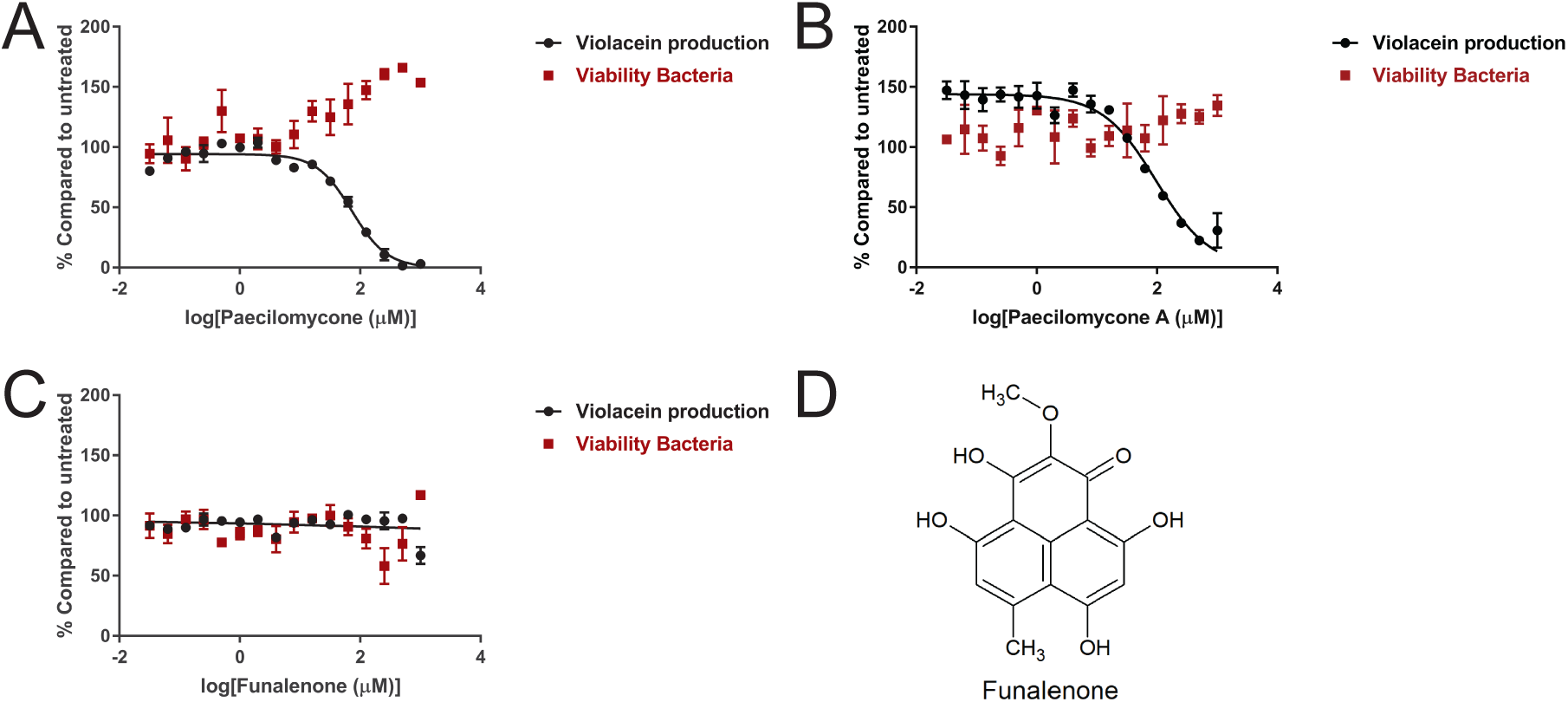
Quorum sensing inhibition by paecilomycone using *C. violaceum* as reporter bacterium. Violacein production and viability of *C. violaceum* after treatment with A) paecilomycone, B) a paecilomycone fraction lacking paecilomycone C, C) Funalenone. D) Structure of funalenone. Experiments were done in triplicates and error bars represent standard error of the mean (SEM).

Whereas the three forms of paecilomycone could not be separated further with the two-step purification we used, we found that, by using TFA instead of ammonium acetate in the second purification step, the amine group was not incorporated in the compound and paecilomycone C was lost from the mixture (Supplementary Figure 5). The fraction lacking paecilomycone C still contained paecilomycone A, a small amount of paecilomycone B, and an unidentified peak. This fraction showed a concentration dependent effect on violacein production with a similar IC_50_ as paecilomycone (IC_50_ = 96.5 µM) (Figure 2B). Toxic effects on the viability of *C. violaceum* were not detected. This suggests that the most abundant paecilomycone A was responsible for most of the QS inhibitor effect of paecilomycone.

Next, we tested the effect of funalenone on QS inhibition. Funalenone is structurally highly related to paecilomycone A with only the position of the methoxy group being different (cf. Figure 2D and 1K). Interestingly, this small difference in structure resulted in a big difference in activity since we did not find inhibition in the production of violacein after funalenone treatment up to a concentration of 1mM (Figure 2C).

Taken together, paecilomycone A and potentially paecilomycone B, but not the highly related funalenone had QS inhibitor activity in *C. violaceum*.

### Paecilomycone shows QS inhibitor activity in *P. aeruginosa*

To test the potential of paecilomycone as QS inhibitor in more clinically relevant bacteria, the effect on various *P. aeruginosa* QS reporters was tested. *P. aeruginosa* PAO1 reporter strains were used that express GFP when the QS pathway is activated, including *lasB-*GFP, *rhlA*-GFP, *pqsA-*GFP. WT-GFP that constitutively expresses GFP was used as control. GFP expression was normalized to the growth of the bacteria and the IC_50_ was calculated by plotting the maximum slope of GFP expression/ bacterial growth (Figure 3).

**Figure 3:**
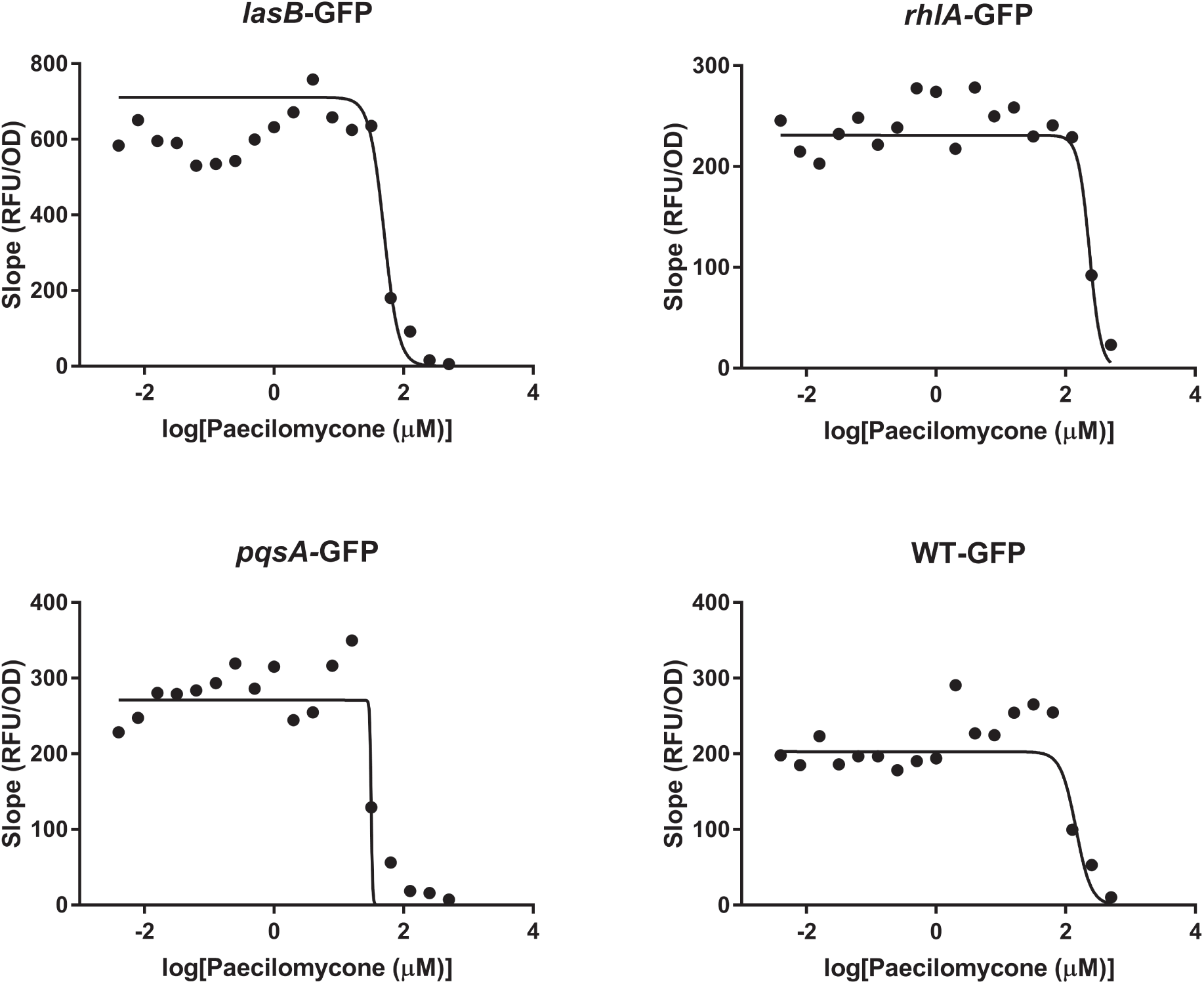
Quorum sensing inhibition in *P. aeruginosa* PAO1 reporter strains after paecilomycone treatment. The effect of paecilomycone was tested using *lasB*-GFP, *rhlA-*GFP, *pqsA-*GFP reporters. In addition, the effect on WT-GFP as control was tested. The maximum slope of RFU, normalized by growth, was plotted and used to calculate the IC_50_. Experiments were done three times in triplicates; the mean of RFU/OD of a representative experiment was plotted.

Paecilomycone inhibited GFP expression in the *lasB-*GFP reporter and *pqsA-* GFP reporter with an IC_50_ of 49.8 µM and 31.5 µM respectively (Figure 3), which is 3- to 4.5-fold lower than in control WT-GFP bacteria (IC_50_ = 143.2 µM), respectively. Concentrations of paecilomycone above 125 µM affected growth of *P. aeruginosa* explaining the effect observed in WT-GFP bacteria (Supplementary Figure 6) .The *rhlA*-GFP reporter was not sensitive to paecilomycone with an apparent IC_50_ of 233.8 µM, which was actually higher than of WT-GFP. We also tested the effect of funalenone on QS inhibition in PAO1. Funalenone did not show an inhibitory effect in any of the reporters (Supplementary Figure 7), indicating that funalenone did not inhibit QS in *P. aeruginosa*.

These results indicate that various, but not all, QS pathways were inhibited by paecilomycone -but not by the highly related funalenone- in clinically relevant *P. aeruginosa* (PAO1) bacteria, with paecilomycone showing the strongest effect in the *pqs*A-GFP reporter.

### Paecilomycone inhibits biofilm formation in *P. aeruginosa*

QS regulates various downstream processes including biofilm formation(27). Therefore, we tested if paecilomycone inhibits biofilm formation by staining biomass using crystal violet after paecilomycone treatment for 24 h. Concentrations of 62.5 µM and higher showed a significant concentration-dependent inhibition of biofilm formation in *P. aeruginosa* PAO1 strain (Figure 4). The highest concentration tested (500 µM) showed an inhibition of 75 % compared to WT control. We conclude that paecilomycone is a potent inhibitor of biofilm formation, because treatment with 62.5 µM already inhibited biofilm formation of *P. aeruginosa* for 50 %.

**Figure 4:**
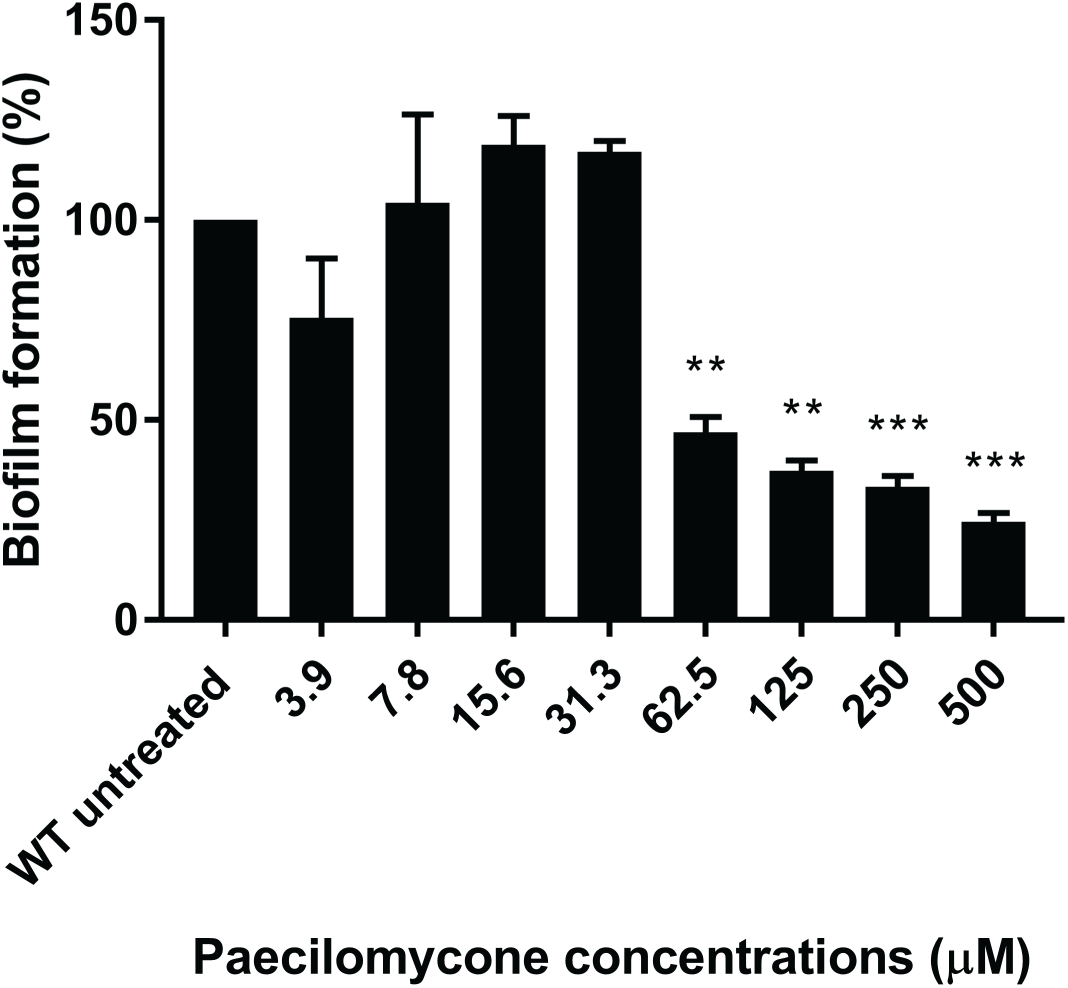
Inhibition of biofilm formation in *P. aeruginosa* PAO1 strain by paecilomycone treatment for 24 h. Biofilm formation was measured by crystal violet staining and normalized to untreated. The mean of three experiments in triplicates was plotted and error bars represent SEM. A one-way ANOVA, corrected for multiple comparisons using Dunnett’s test was done to determine statistical significance. Treated samples were compared to untreated controls (**, P<0.005; ***, P<0.001)

### Paecilomycone alters the production of various virulence factors

Besides biofilm formation, QS regulates the synthesis of virulence factors in *P. aeruginosa*(28–30). Therefore, we tested the effect of paecilomycone treatment on the synthesis of various virulence factors. Paecilomycone induced a strong and significant inhibitory effect on the production of pyocyanin with a maximum inhibition of 88 % and an IC_50_ of 8.5 µM (Figure 5A). The lowest concentration tested, 2.0 µM, still showed a significant decrease in pyocyanin production. To rule out inadvertent effects on PAO1 due to altered growth, growth kinetics were determined in King’s A medium, the medium used for analysis of pyocyanin. The initial slopes of the PAO1 bacterial density curves were similar in control and paecilomycone treated samples, indicating that bacterial growth was similar (Supplementary Figure 8). Yet, bacterial density did not reach the same level in samples treated with high paecilomycone concentrations (from 31.3 µM onwards). However, the difference in bacterial densities after 24 h treatment was relatively small and could not explain the large decrease in pyocyanin production in response to paecilomycone.

**Figure 5:**
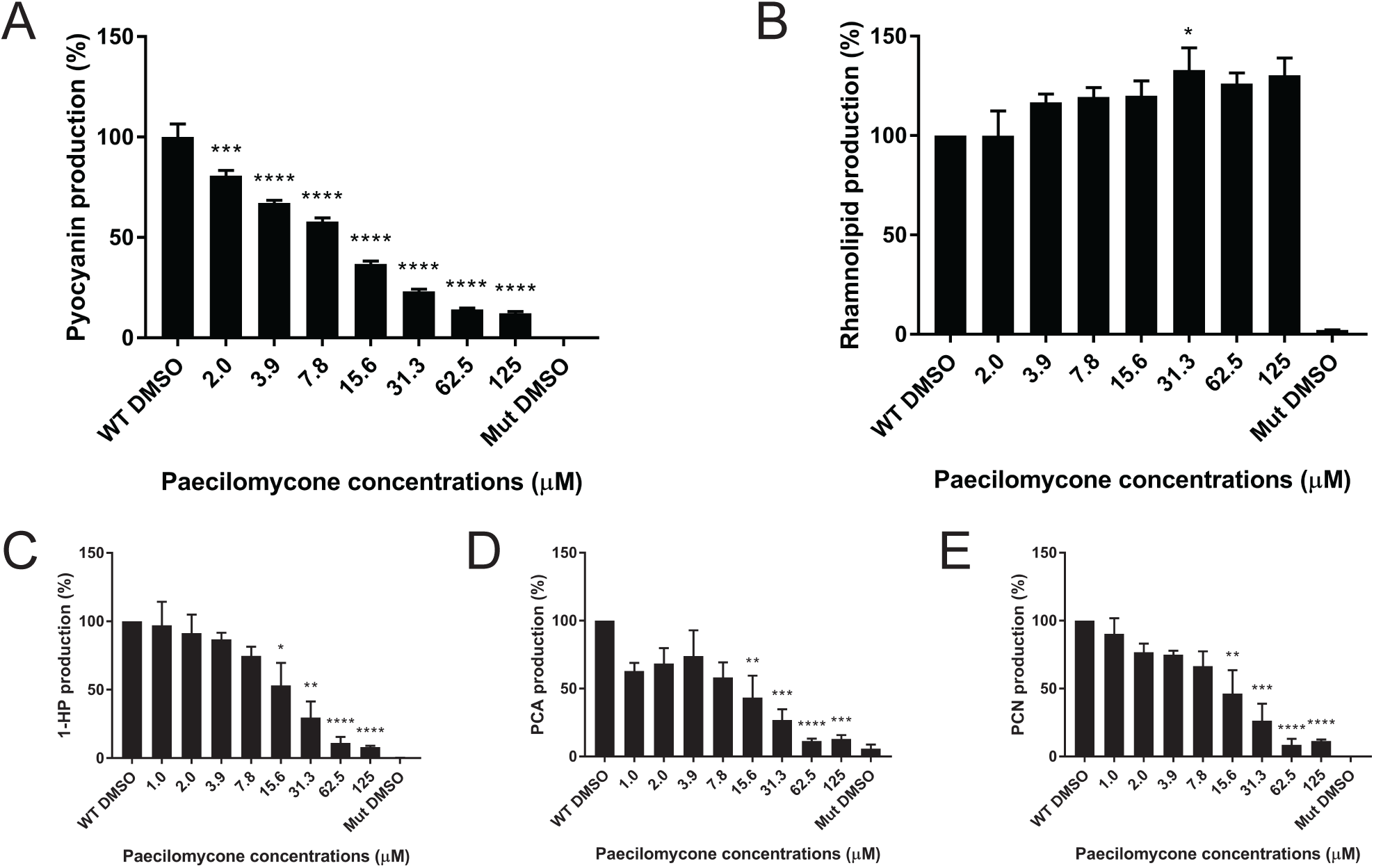
Production of virulence factors by *P. aeruginosa* PAO1 strain after paecilomycone treatment. Graphs show the production of A) pyocyanin, B) rhamnolipids, C) 1-hydroxyphenazine (1-HP), D) phenazine-1-carboxylic acid (PCA), E) phenazine-1-carboxamide (PCN), after treatment with paecilomycone for 24 h. A QS mutant (Δ*lasI*/Δ*rhlI*) was included as negative control. Experiments were performed in biological triplicates containing technical triplicates (with exception of pyocyanin, which was performed once in this setting). Values were normalized to WT DMSO control and the mean was plotted, error bars represent SEM. A one-way ANOVA, corrected for multiple comparisons using Dunnett’s test was done to determine statistical significance. Paecilomycone treated samples were compared to WT DMSO control (*, P<0.05; **, P<0.005; ***, P<0.001; ****, P<0.0001).

Interestingly, rhamnolipid production in PAO1 cells in response to paecilomycone was increased (Figure 5B). A modest but significant increase, up to 33 %, was observed following treatment with 31 µM paecilomycone.

Because the synthesis of pyocyanin showed a strong inhibition, we also measured the effect of paecilomycone on other phenazines produced by *P. aeruginosa*: 1-hydroxyphenazine (1-HP), phenazine-1-carboxylic acid (PCA), and phenazine-1-carboxamide (PCN)(28). The synthesis of these phenazines all showed a significant inhibition after paecilomycone treatment with a concentration of 15.6 µM or higher (Figure 5C-E). Maximum inhibition was reached after treatment with 62.5- 125 µM paecilomycone with inhibition up to 92 % (1-HP), 89 % (PCA), and 91 % (PCN) compared to untreated. This shows that paecilomycone inhibited the production of phenazines, but not rhamnolipids.

### Paecilomycone inhibits production of PQS pathway metabolites

Since the strongest inhibition of QS was measured in the PQS pathway (Figure 3) and a strong inhibition was observed in the synthesis of phenazines (Figure 5), we tested the effect of paecilomycone treatment on the production of various metabolites in PQS synthesis (Figure 6A). In short, PQS synthesis starts with PqsA, which converts anthranilic acid to anthraniloyl-CoA(31–33). After condensation by PqsD, PqsE hydrolyses 2-aminobenzoylacetyl-CoA (2-ABA-CoA) into 2-aminobenzoylacetyl (2-ABA)(34–36). This hydrolysis can be taken over by TesB thioesterase(36). 2-ABA is condensed to HHQ by the PqsBC complex which is hydroxylated by PqsH to form PQS(34, 37). In addition, 2-ABA can spontaneously decarboxylate to 2’- aminoacetophenone (2-AA), or form 2-heptyl-4-quinolinol 1-oxide (HQNO) by PqsL and the PqsBC complex(11, 34, 38). To test the effect of paecilomycone on PQS synthesis, we tested the production of 2-AA, HQNO, HHQ, and PQS.

**Figure 6:**
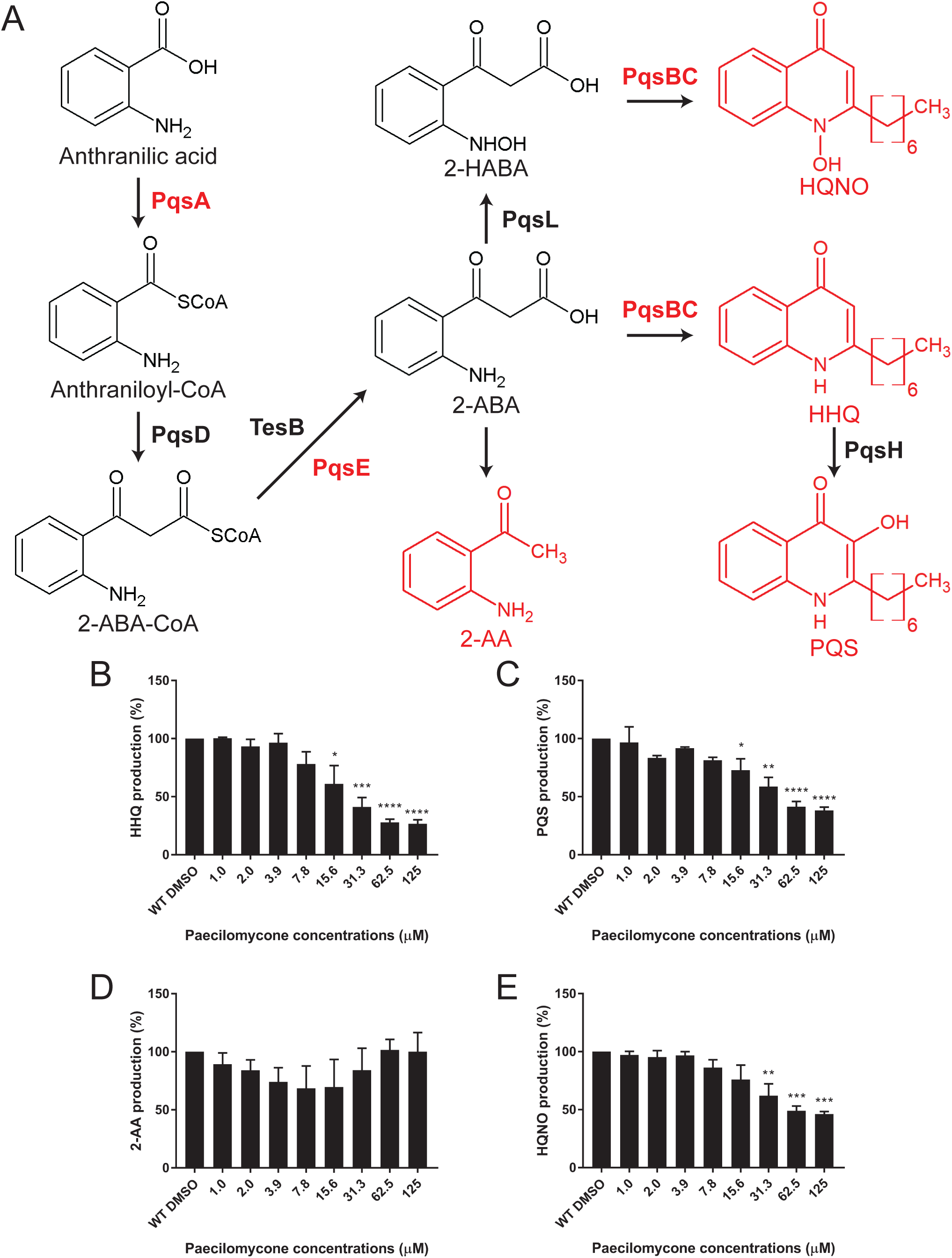
Inhibition of various metabolites in PQS synthesis after paecilomycone treatment. A) PQS synthesis pathway starts from anthranilic acid which is converted by enzymes from the *pqsABCDE* operon to produce HHQ, which is then hydroxylated by PqsH to form PQS. 2-ABA can also spontaneously decarboxylate into 2-AA or be converted into HQNO by PqsL and subsequently the PqsBC complex. Enzymes in red indicate mutants used in figure 7. Graphs show the production of the metabolites indicated in red: B) HHQ, C) PQS, D) 2-AA, E) HQNO. Experiments were done three times in triplicates, values were normalized to DMSO treated control, and the mean of the experiments is plotted with error bars representing the SEM. A one-way ANOVA, corrected for multiple comparisons using Dunnett’s test, was done to determine statistical significance. Treated samples were compared to DMSO control (*, P<0.05; **, P<0.005; ***, P<0.001; ****, P<0.0001).

After treatment for 24 h, we measured a significant concentration-dependent inhibition of HHQ and PQS using paecilomycone concentrations of 15.6 µM and higher (Figure 6B-C). The strongest inhibition was measured after treatment with 125 µM paecilomycone with inhibition up to 73 % and 62 % for HHQ and PQS, respectively. Besides HHQ and PQS production, paecilomycone also significantly inhibited HQNO production up to 54 % compared to control (Figure 6D). Interestingly, the production of 2-AA was not significantly affected after paecilomycone treatment (Figure 6E). This suggests that paecilomycone treatment did not inhibit the complete PQS synthesis, but only specific enzymes like the PqsBC complex.

To exclude the possibility that paecilomycone C inhibited phenazine production and the PQS pathway, we compared the activity of paecilomycone with the fraction without paecilomycone C. We observed no differences, indicating that paecilomycone C is dispensable for inhibition of phenazine production (Supplementary figure 9).

### Paecilomycone is a potential inhibitor of the PqsBC complex

Since various metabolites of the PQS pathway were inhibited, except for 2-AA, we expected paecilomycone to target the PqsBC complex (Figure 6A). PqsBC inhibitors have been reported before and show similar patterns as paecilomycone treatment(39–41). To test involvement of the *pqs* genes in the PQS synthesis pathway in our conditions (King’s A medium), we analyzed metabolite expression in PAO1 mutants, which lack genes encoding PQS synthesis enzymes (Δ*pqsA,* Δ*pqsE,* Δ*pqsBC*). We expected the *pqsBC* mutant to show a similar pattern as paecilomycone treatment, *i.e.* inhibition of HQNO, HHQ, PQS, pyocyanin and PCA production while not affecting 2-AA.

As expected, we measured a strong inhibition of both HHQ and PQS in Δ*pqsA* and Δ*pqsBC* (Figure 7A,B). Δ*pqsE* only showed a slight decrease in PQS and even an increase in HHQ (Figure 7A-B), which may be due to TesB, which hydrolyses 2- ABA-CoA to 2-ABA in absence of PqsE(36) (Figure 6A). HQNO production was significantly inhibited in all mutant strains, with complete inhibition in both *pqsA* and *pqsBC* mutants (Figure 7C). In contrast to paecilomycone treatment, which did not affect 2-AA, the *pqsA* mutant showed almost a 100% decrease in 2-AA production, which was expected because PqsA mediates the first step in PQS synthesis (Figure 7D). Surprisingly, the *pqsBC* mutant still showed a strong decrease in 2-AA production. This may be due to inhibition of the positive feedback of HHQ and PQS on the *pqsABCDE* operon upon complete inhibition of HHQ and PQS production(32).

**Figure 7:**
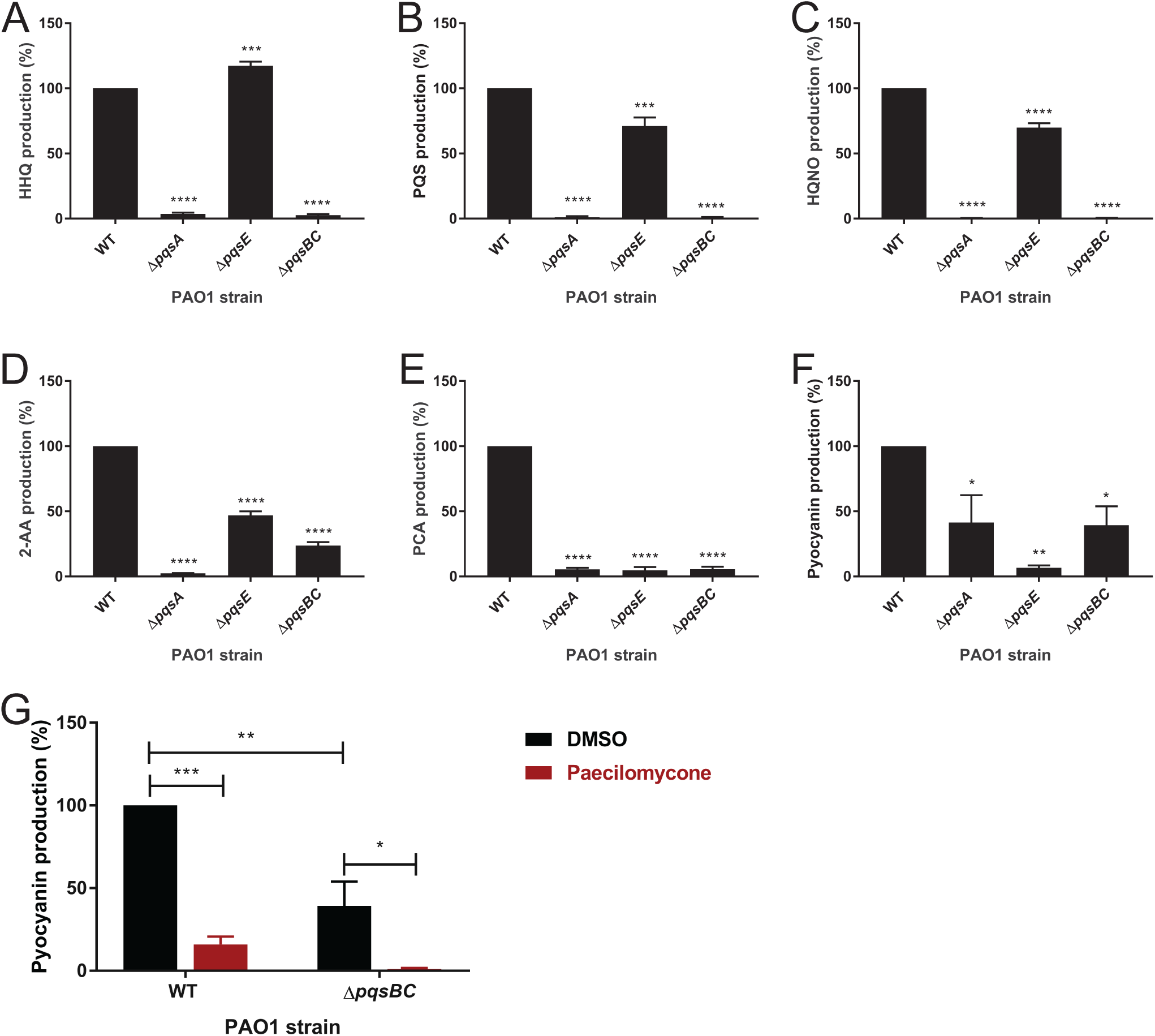
The effect of mutations in PQS synthesis enzymes on the production of various HAQs, phenazines and 2-AA in King’s A medium. *P. aeruginosa* PAO1 strain and various mutant (Δ*pqsA* / Δ*pqsE* / Δ*pqsBC*) were grown in King’s A medium for 24 h and the production of A) HHQ, B) PQS, C) HQNO, D) 2-AA, E) PCA, F) pyocyanin was measured and normalized to WT control. G) WT and *ΔpqsBC* mutants were treated with paecilomycone and pyocyanin levels were measured. Experiments were done three times in triplicates, the mean of the normalized values is plotted with error bars representing the SEM. A one-way ANOVA, corrected for multiple comparisons using Dunnett’s test, was done to determine statistical significance. Mutants were compared to WT control (*, P<0.05; **, P<0.005; ***, P<0.001; ****, P<0.0001).

PqsE is responsible for the synthesis of pyocyanin. Consistent with this notion, production of pyocyanin and PCA was abolished in *pqsE* mutant. Production of PCA and pyocyanin was also abolished or reduced in *pqsA* and *pqsBC* mutants (Figure 7E- F). This may be due to positive feedback, leading to reduced expression of PqsE in *pqsA* and *pqsBC* mutants. Together, these data suggest that paecilomycone treatment did not completely inhibit the PQS pathway (*cf.* Δ*pqsA*), or specifically target phenazine synthesis (*cf.* Δ*pqsE*), but rather that paecilomycone inhibited aspects of PQS synthesis (*cf.* Δ*pqsBC*).

Interestingly, while the *pqsBC* mutant already showed decreased levels of pyocyanin, treatment with paecilomycone strengthened this inhibition significantly to almost 100% (Figure 7G). This suggests that paecilomycone might have alternative targets, or targeted a process related to the enzymatic activity of the PqsBC complex, which caused an even stronger inhibition of pyocyanin in the absence of the PqsBC complex.

### Toxicity of paecilomycone

To test if paecilomycone has clinical potential, we determined the cytotoxicity of paecilomycone on various viability models. First, we used human liver-derived HepG2 cells. Viability of HepG2 cells was assessed after paecilomycone treatment for 24 h. Paecilomycone was toxic to HepG2 cells at high concentrations with an IC_50_ of 219 µM (Figure 8A). However, up to a concentration of 125 µM paecilomycone, there was no difference in the viability of HepG2 cells compared to untreated control cells.

**Figure 8:**
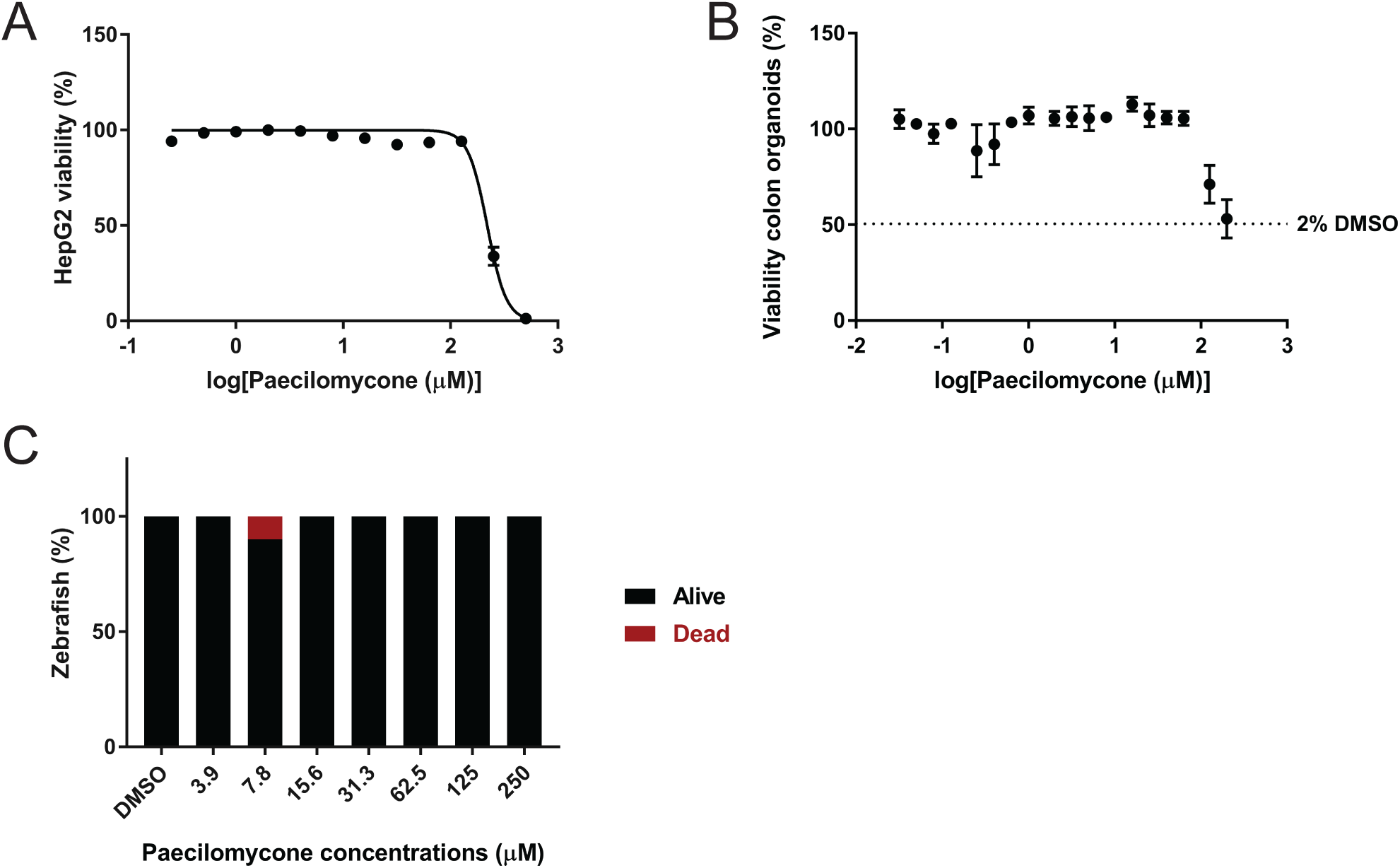
Toxicity of paecilomycone treatment using various toxicity models. A) Viability of HepG2 cells treated with paecilomycone for 24 h, viability was measured using resazurin assay. B) Viability of colon organoids treated with paecilomycone for 5 d, viability was measured by measuring ATP levels by a Cell Titer GLO 3D assay. C) Viability of zebrafish embryos that were treated after 48 h for 24h.

Second, we tested the cytotoxic effect of paecilomycone on more complex systems using human colon organoids. Viability of the organoids was measured after paecilomycone treatment for 5 days. High concentrations (200 µM and 120 µM) showed reduced viability compared to 1% DMSO (Figure 8B). However, at these high concentrations, the final concentration of paecilomycone solvent was higher than 1%, up to 2 % DMSO, which by itself already showed a 50 % reduction in viability. It is therefore highly likely that the apparent effect of 120 µM and 200 µM paecilomycone on organoid viability was actually caused by high concentrations of DMSO.

Third, we tested the cytotoxic effect of paecilomycone on a whole organism *in vivo*, 2-day old zebrafish embryos. The viability of the embryos was measured after paecilomycone treatment for 24 h. Each concentration was tested on approximately 10 embryos. No toxic effect of paecilomycone was apparent (Figure 8C).

In conclusion, paecilomycone did not affect eukaryotic cells at concentrations that inhibited QS in Gram-negative bacteria. Only at high paecilomycone concentrations, an effect was detected on HepG2 cells, whereas no effect was specifically attributable to paecilomycone treatment of more complex systems, like organoids and zebrafish embryos.

## Discussion

In this study, we describe the identification of paecilomycone as a novel QS inhibitor. Paecilomycone showed strong inhibition of QS in Gram-negative bacterial strains, *C. violaceum* and *P. aeruginosa* PAO1. Paecilomycone mainly inhibited the PQS pathway and it strongly inhibited the production of HHQ and PQS. In addition, paecilomycone inhibited biofilm formation and the production of phenazines, including pyocyanin. Paecilomycone might have promising clinical value because cytotoxicity was very low.

Paecilomycone treatment showed strong inhibition of QS in the *pqsA*-GFP reporter (Figure 3), production of related virulence factors (Figure 5), and production of HHQ and PQS (Figure 6). Since paecilomycone did not interfere with the production of 2-AA (Figure 6), we suggested that paecilomycone affected the activity of the PqsBC complex. Only a limited number of PqsBC inhibitors are identified so far. Previously described PqsBC inhibitors show similar profiles as paecilomycone: inhibition of HHQ and PQS while 2-AA is unaffected or even upregulated(39–41). Only dual PqsBC / PqsR inhibitors showed inhibition of 2-AA, something that is comparable with our Δ*pqsBC* mutant strain(39–41). Unfortunately, these studies do not describe the effect of the PqsBC inhibitors on expression of virulence traits in *P. aeruginosa* except for a small decrease in pyocyanin production, whereas we observed a strong inhibition after paecilomycone treatment(39). Based on our data, the PqsBC complex is the most likely target of paecilomycone treatment.

PqsBC complex might be a direct target of paecilomycone. However, PqsE instead of PQS controls the production of phenazines via RhlR(42–47). A mutant lacking PqsE did not produce PCA or pyocyanin, underlining the importance of PqsE for production of these phenazines. The observed inhibition of PCA and pyocyanin production in Δ*pqsA* and Δ*pqsBC* may be explained by inhibition of the *pqsABCDE* operon via inhibition of a feed-forward loop, which eventually results in reduced expression of PqsE. However, paecilomycone treatment does not inhibit 2-AA production; therefore, we do not expect a strong inhibition of the *pqsABCDE* operon by paecilomycone treatment. Moreover, paecilomycone still had a significant inhibitory effect on pyocyanin production in the Δ*pqsBC* strain (Figure 7G), suggesting that the PqsBC complex cannot be the only target of paecilomycone. Because of the strong inhibition of phenazines, and inhibition of pyocyanin in the *pqsBC* mutant, we believe that paecilomycone did not only target the PqsBC complex. Instead, we suggest that paecilomycone affects QS by directly targeting the PqsBC complex and alternative targets, or paecilomycone alters processes that influence the enzymatic activity of the PqsBC complex. Further research will be needed to elucidate the exact mechanism of action of paecilomycone.

Alternative targets or alteration of upstream processes could also explain the inhibition in QS in both *C. violaceum* (Figure 2) and *P. aeruginosa* (Figure 3). Although both bacteria contain LuxI/R-type QS, the PQS system is unique for *P. aeruginosa* (9–11, 25). More specifically, the PqsBC complex is a FabH-like enzyme that catalyzes the condensation of an octanoyl-CoA with 2-ABA(48, 49). This complex does not show similarity with the CviI synthase in *C. violaceum*. Therefore, interaction of paecilomycone with a LuxI/R type system of *P. aeruginosa* would be expected. For example, Welsh *et al* have discovered RhlR agonists that also show strong inhibition of pyocyanin production by inhibition of the PQS system by RhlR activation(50). However, this activation would also lead to inhibition of 2-AA since the complete *pqsABCDE* operon will be inhibited. Therefore, we believe paecilomycone has a different mechanism of action in *C. violaceum*.

Paecilomycone consisted of three different paecilomycones. Analytical HPLC analysis showed that the concentration of paecilomycone A is highest (Figure 1). To validate that paecilomycone A is responsible for the measured activity, we tested the purified fraction without paecilomycone C (Supplementary Figure 5) in various assays. Both in *C. violaceum* (Figure 2) and in the inhibition of phenazines and HAQs (Supplementary Figure 9) we measured similar activities between paecilomycone and the fraction lacking paecilomycone C, suggesting that paecilomycone C was not required for the effect of paecilomycone in QS inhibition. Interestingly, although funalenone shows high resemblance with paecilomycone A, funalenone does not show inhibition of QS in *C. violaceum* (Figure 2) or *P. aeruginosa* (Supplementary Figure 7). This suggests that the position of the methoxy group is important in the QS inhibitor activity.

Inhibition of HAQ and phenazine production is interesting to fight the virulence of *P. aeruginosa*. PQS is important in the upregulation of various genes involved in *P. aeruginosa* virulence including the Type III-secretion system, ExoS toxin, and siderophore synthesis(42). Phenazines are one of the major virulence factors produced by *P. aeruginosa,* and play an important role in the acute and chronic lung infections through various mechanisms(51–55). Phenazines have been shown to be toxic in various model organisms, including *C. elegans*(56), *Drosophila melanogaster*(57), mice(58) and epithelial lung cells(54, 55). In addition, in humans, phenazine production has been measured in sputum samples in concentrations that are able to inhibit ciliary beating, which is important for the clearance of bacteria(59). Moreover, phenazines are also involved in biofilm formation(60) and in the acquisition of iron(61). In general, PQS and phenazines are important for the survival of the bacteria and for damaging the host. Therefore, inhibition of PQS and phenazines by paecilomycone holds the potential as interesting therapeutic lead, by reducing toxicity and subsequently morbidity and mortality in patients.

Because of its potential to inhibit QS and downstream targets like phenazine production and biofilm formation, we believe paecilomycone is an interesting lead for further clinical research. We showed that concentrations that inhibit virulence factor production and biofilm formation are not toxic to cell cultures. Moreover, we did not measure toxic effects in more complex systems up to the highest concentrations tested (200-250 µM). This makes paecilomycone an interesting lead for further research as potential agent to fight *P. aeruginosa* infections.

## Author Contributions

WAGB and JdH conceived and designed the study. WAGB wrote the first draft of the manuscript. WAGB, JH and MBH performed experiments. JdH supervised the project, acquired funding and administered the project. All authors contributed to manuscript revision, read, and approved the submitted version.

## Acknowledgments

We would like to thank Tim Holm Jakobsen, and Bart W. Bardoel for sharing bacterial strains with us. We would like to thank Joe J. Harrison for sharing the pEX18Gm plasmid. We thank Justyna M. Dobruchowska and Geert-Jan Boons for help with NMR and Arjan Barendregt and Albert Heck for help with high-resolution mass spectrometry.

## References

1. Botelho, J., Grosso, F. & Peixe, L. Antibiotic resistance in *Pseudomonas aeruginosa* – Mechanisms, epidemiology and evolution. Drug Resist. Updat. 44, 100640 (2019).

2. Moore, N. M. & Flaws, M. Introduction: *Pseudomonas aeruginosa*. Clin. Lab. Sci. 24, 41–42 (2011).

3. Burrows, L. L. The Therapeutic Pipeline for *Pseudomonas aeruginosa* Infections. ACS Infect. Dis. 4, 1041–1047 (2018).

4. Parkins, M. D., Somayaji, R. & Waters, V. J. Epidemiology, biology, and impact of clonal pseudomonas aeruginosa infections in cystic fibrosis. Clin. Microbiol. Rev. 31, (2018).

5. Williams, H. D. & Davies, J. C. Basic science for the chest physician: *Pseudomonas aeruginosa* and the cystic fibrosis airway. Thorax 67, 465–467 (2012).

6. Bassler, B. L. & Losick, R. Bacterially Speaking. Cell 125, 237–246 (2006).

7. Defoirdt, T. Quorum-Sensing Systems as Targets for Antivirulence Therapy. Trends Microbiol. 26, 313–328 (2018).

8. Whitehead, N. A., Barnard, A. M. L., Slater, H., Simpson, N. J. L. & Salmond, G. P. C. Quorum-sensing in Gram-negative bacteria. FEMS Microbiol. Lett. 25, 365–404 (2001).

9. Kostylev, M. et al. Evolution of the *Pseudomonas aeruginosa* quorum-sensing hierarchy. Proc. Natl. Acad. Sci. U S A 116, 7027–7032 (2019).

10. Lee, J. & Zhang, L. The hierarchy quorum sensing network in *Pseudomonas aeruginosa*. Protein Cell 6, 26–41 (2014).

11. García-Reyes, S., Soberón-Chávez, G. & Cocotl-Yanez, M. The third quorum--sensing system of *Pseudomonas aeruginosa*: *Pseudomonas* quinolone signal and the enigmatic PqsE protein. J. Med. Microbiol. 69, 25–34 (2020).

12. Soukarieh, F., Williams, P., Stocks, M. J. & Cámara, M. *Pseudomonas aeruginosa* Quorum Sensing Systems as Drug Discovery Targets: Current Position and Future Perspectives. J. Med. Chem. 61, 10385–10402 (2018).

13. Schütz, C. & Empting, M. Targeting the *Pseudomonas* quinolone signal quorum sensing system for the discovery of novel anti-infective pathoblockers. Beilstein J. Org. Chem. 14, 2627–2645 (2018).

14. Jakobsen, T. H. et al. Ajoene, a Sulfur-Rich Molecule from Garlic, Inhibits Genes Controlled by Quorum Sensing. Antimicrob. Agents Chemother. 56, 2314–2325 (2012).

15. D’Angelo, F. et al. Identification of FDA-approved drugs as antivirulence agents targeting the pqs Quorum-Sensing system of *Pseudomoans aeruginosa*. Antimicrob. Agents Chemother. 62, 1–20 (2018).

16. Beenker, W. A. G., Hoeksma, J. & Hertog, J. Den. Gregatins, a Group of Related Fungal Secondary Metabolites, Inhibit Aspects of Quorum Sensing in Gram- Negative Bacteria. Front. Microbiol. 13, (2022).

17. Hentzer, M. & Givskov, M. Pharmacological inhibition of quorum sensing for the treatment of chronic bacterial infections. J. Clin. Invest. 112, 1300–1307 (2003).

18. Yang, L. et al. Effects of iron on DNA release and biofilm development by *Pseudomonas aeruginosa*. Microbiology 153, 1318–1328 (2007).

19. Hentzer, M. et al. Inhibition of quorum sensing in *Pseudomonas aeruginosa* biofilm bacteria by a halogenated furanone compound. Microbiology 148, 87– 102 (2002).

20. Fong, J. et al. Disulfide Bond-Containing Ajoene Analogues As Novel Quorum Sensing Inhibitors of *Pseudomonas aeruginosa*. J. Med. Chem. 60, 215–227 (2017).

21. Hentzer, M. et al. Attenuation of *Pseudomonas aeruginosa* virulence by quorum sensing inhibitors. EMBO J. 22, 3803–3815 (2003).

22. Hmelo, L. R. et al. Precision-engineering the *Pseudomonas aeruginosa* genome with two-step allelic exchange. Nat. Protoc. 10, 1820–1841 (2015).

23. Skogman, M. E., Kanerva, S., Manner, S., Vuorela, P. M. & Fallarero, A. Flavones as quorum sensing inhibitors identified by a newly optimized screening platform using *Chromobacterium violaceum* as reporter bacteria. Molecules 21, (2016).

24. Jakobsen, T. H., Alhede, M., Hultqvist, L. D., Bjarnsholt, T. & Givskov, M. Qualitative and Quantitative Determination of Quorum Sensing Inhibition In Vitro. Quor. Sens. Methods Protoc. 1673, 275–285 (2010).

25. Essar, D. W., Eberly, L., Hadero, A. & Crawford, I. P. Identification and characterization of genes for a second anthranilate synthase in *Pseudomonas aeruginosa*: Interchangeability of the two anthranilate synthase and evolutionary implications. J. Bacteriol. 172, 884–900 (1990).

26. Zhou, S., Zhang, A. & Chu, W. Phillyrin is an effective inhibitor of quorum sensing with potential as an anti-*Pseudomonas aeruginosa* infection therapy. J. Vet. Med. Sci. 81, 473–479 (2019).

27. Pleguezuelos-Manzano, C. et al. Establishment and Culture of Human Intestinal Organoids Derived from Adult Stem Cells. Curr. Protoc. Immunol. 130, (2020).

28. Driehuis, E., Kretzschmar, K. & Clevers, H. Establishment of patient-derived cancer organoids for drug-screening applications. Nat. Protoc. 15, 3380–3409 (2020).

29. Aleström, P., et al. Zebrafish: Housing and husbandry recommendations. Lab. Anim. 54, 213–224 (2020).

30. Stauff, D. L., Bassler, B. L., Hughes, H. & Chase, C. Quorum Sensing in *Chromobacterium violaceum* : DNA Recognition and Gene Regulation by the CviR Receptor. J. Bacteriol. 193, 3871–3878 (2011).

31. Lu, R. et al. New tyrosinase inhibitors from *Paecilomyces gunnii*. J. Agric. Food Chem. 62, 11917–11923 (2014).

32. Jakobsen, T. H., Tolker-Nielsen, T. & Givskov, M. Bacterial biofilm control by perturbation of bacterial signaling processes. Int. J. Mol. Sci. 18, (2017).

33. Nadal Jimenez, P., et al. The Multiple Signaling Systems Regulating Virulence in *Pseudomonas aeruginosa*. Microbiol. Mol. Biol. Rev. 76, 46–65 (2012).

34. Brint, J. M. & Ohman, D. E. Synthesis of multiple exoproducts in *Pseudomonas aeruginosa* is under the control of RhlR-RhlI, another set of regulators in strain PAO1 with homology to the autoinducer-responsive LuxR-LuxI family. J. Bacteriol. 177, 7155–7163 (1995).

35. Ochsner, U. A. & Reiser, J. Autoinducer-mediated regulation of rhamnolipid biosurfactant synthesis in *Pseudomonas aeruginosa*. Proc. Natl. Acad. Sci. U. S. A. 92, 6424–6428 (1995).

36. Calfee, M. W., Coleman, J. P. & Pesci, E. C. Interference with *Pseudomonas* quinoline signal synthesis inhibits virulence factor expression by *Pseudomonas aeruginosa*. Proc. Natl. Acad. Sci. U. S. A. 98, 11633–11637 (2001).

37. Déziel, E. et al. Analysis of *Pseudomonas aeruginosa* 4-hydroxy-2- alkylquinolines (HAQs) reveals a role for 4-hydroxy-2-heptylquinoline in cell-to- cell communication. Proc. Natl. Acad. Sci. U. S. A. 101, 1339–1344 (2004).

38. Coleman, J. P. et al. *Pseudomonas aeruginosa* PqsA is an anthranilate- coenzyme A ligase. J. Bacteriol. 190, 1247–1255 (2008).

39. Dulcey, C. E. et al. The end of a long-standing hypothesis: the *Pseudomonas* signalling molecules 4-hydroxy-2-alkylquinolines are derived from fatty acids, not 3-ketofatty acids. Chem. Biol. 20, (2013).

40. Zhang, Y. M., Frank, M. W., Zhu, K., Mayasundari, A. & Rock, C. O. PqsD is responsible for the synthesis of 2,4-dihydroxyquinoline, an extracellular metabolite produced by *Pseudomonas aeruginosa*. J. Biol. Chem. 283, 28788– 28794 (2008).

41. Drees, S. L. & Fetzner, S. PqsE of *Pseudomonas aeruginosa* acts as pathway- specific thioesterase in the biosynthesis of alkylquinolone signaling molecules. Chem. Biol. 22, 611–618 (2015).

42. Schertzer, J. W., Brown, S. A. & Whiteley, M. Oxygen Levels Rapidly Modulate *Pseudomonas aeruginosa* Social Behaviors via Substrate Limitation of PqsH. Mol. Microbiol. 77, 1527–1538 (2010).

43. Drees, S. L. et al. PqsL uses reduced flavin to produce 2- hydroxylaminobenzoylacetate, a preferred PqsBC substrate in alkyl quinolone biosynthesis in *Pseudomonas aeruginosa*. J. Biol. Chem. 293, 9345–9357 (2018).

44. Starkey, M. et al. Identification of Anti-virulence Compounds That Disrupt Quorum-Sensing Regulated Acute and Persistent Pathogenicity. PLoS Pathog. 10, (2014).

45. Allegretta, G. et al. In-depth profiling of MvfR-regulated small molecules in *Pseudomonas aeruginosa* after Quorum Sensing inhibitor treatment. Front. Microbiol. 8, 1–12 (2017).

46. Maura, D. et al. Polypharmacology Approaches against the *Pseudomonas aeruginosa* MvfR Regulon and Their Application in Blocking Virulence and Antibiotic Tolerance. ACS Chem. Biol. 12, 1435–1443 (2017).

47. Rampioni, G. et al. Unravelling the Genome-Wide Contributions of Specific 2- Alkyl-4-Quinolones and PqsE to Quorum Sensing in *Pseudomonas aeruginosa*. PLoS Pathog. 12, 1–25 (2016).

48. Déziel, E. et al. The contribution of MvfR to *Pseudomonas aeruginosa* pathogenesis and quorum sensing circuitry regulation: Multiple quorum sensing- regulated genes are modulated without affecting IasRI, rhIRI or the production of N-acyl-L-homoserine lactones. Mol. Microbiol. 55, 998–1014 (2005).

49. Farrow, J. M. et al. PqsE functions independently of PqsR-*Pseudomonas* quinolone signal and enhances the rhl quorum-sensing system. J. Bacteriol. 190, 7043–7051 (2008).

50. Hazan, R. et al. Homeostatic interplay between bacterial cell-cell signaling and iron in virulence. PLoS Pathog. 6, (2010).

51. Mukherjee, S. et al. The PqsE and RhlR proteins are an autoinducer synthase– receptor pair that control virulence and biofilm development in *Pseudomonas aeruginosa*. Proc. Natl. Acad. Sci. U. S. A. 115, E9411–E9418 (2018).

52. Simanek, K. A., Taylor, I. R., Richael, E. K. & Lasek-nesselquist, E. The PqsE- RhlR Interaction Regulates RhlR DNA Binding to Control Virulence Factor Production in *Pseudomonas aeruginosa*. Microbiol. Spectr. 10, (2022).

53. Witzgall, F. et al. The alkylquinolone repertoire of *Pseudomonas aeruginosa* is linked to structural flexibility of the fabh-like 2-heptyl-3-hydroxy-4(1h)-quinolone (pqs) biosynthesis enzyme pqsbc. ChemBioChem 19, 1531–1544 (2018).

54. Drees, S. L. et al. PqsBC, a condensing enzyme in the biosynthesis of the pseudomonas aeruginosa quinolone signal: Crystal structure, inhibition, and reaction mechanism. J. Biol. Chem. 291, 6610–6624 (2016).

55. Welsh, M. A., Eibergen, N. R., Moore, J. D. & Blackwell, H. E. Small Molecule Disruption of Quorum Sensing Cross-Regualtion in *Pseudomonas aeruginosa* Causes Major and Unexpected Alterations to Virulence Phenotypes. J. Am. Chem. Soc. 137, 1510–1519 (2015).

56. Vilaplana, L. & Marco, M. P. Phenazines as potential biomarkers of *Pseudomonas aeruginosa* infections: synthesis regulation, pathogenesis and analytical methods for their detection. Anal. Bioanal. Chem. 412, 5897–5912 (2020).

57. Lau, G. W., Hassett, D. J., Ran, H. & Kong, F. The role of pyocyanin in *Pseudomonas aeruginosa* infection. Trends Mol. Med. 10, 599–606 (2004).

58. Price-Whelan, A., Dietrich, L. E. P. & Newman, D. K. Rethinking ‘secondary’ metabolism: Physiological roles for phenazine antibiotics. Nat. Chem. Biol. 2, 71–78 (2006).

59. Hall, S. et al. Cellular effects of pyocyanin, a secreted virulence factor of *Pseudomonas aeruginosa*. Toxins (Basel). 8, 1–14 (2016).

60. Rada, B. & Leto, T. L. Pyocyanin effects on respiratory epithelium: Relevance in *Pseudomonas aeruginosa* airway infections. Trends Microbiol. 21, 73–81 (2013).

61. Mahajan-Miklos, S., Tan, M. W., Rahme, L. G. & Ausubel, F. M. Molecular mechanisms of bacterial virulence elucidated using a *Pseudomonas aeruginosa*-*Caenorhabditis elegans* pathogenesis model. Cell 96, 47–56 (1999).

62. Lau, G. W. et al. The *Drosophila melanogaster* toll pathway participates in resistance to infection by the gram-negative human pathogen *Pseudomonas aeruginosa*. Infect. Immun. 71, 4059–4066 (2003).

63. Lau, G. W., Ran, H., Kong, F., Hassett, D. J. & Mavrodi, D. *Pseudomonas aeruginosa* pyocyanin is critical for lung infection in mice. Infect. Immun. 72, 4275–4278 (2004).

64. Wilson, R. et al. Measurement of *Pseudomonas aeruginosa* phenazine pigments in sputum and assessment of their contribution to sputum sol toxicity for respiratory epithelium. Infect. Immun. 56, 2515–2517 (1988).

65. Ramos, I., Dietrich, L. E. P., Price-Whelan, A. & Newman, D. K. Phenazines affect biofilm formation by *Pseudomonas aeruginosa* in similar ways at various scales. Res. Microbiol. 161, 187–191 (2010).

66. Cox, C. D. Role of pyocyanin in the acquisition of iron from transferrin. Infect. Immun. 52, 263–270 (1986).

